# Nirmatrelvir-Resistant Mutations in SARS-CoV-2 Mpro Enhance Host Immune Evasion via Cleavage of NF-κB Essential Modulator

**DOI:** 10.1101/2024.10.18.619137

**Authors:** Merrilee Thomas, Thom Hughes

## Abstract

Nirmatrelvir is a SARS-CoV-2 M^pro^ inhibitor in Paxlovid. Patients treated with it often produce mutant viruses in which the M^pro^ resists Nirmatrelvir inhibition. A common interpretation is that the mutations allow the virus to escape inhibition, but here we report that these mutations enable the protease to more effectively cleave the host protein NF-kappa-B essential modulator (NEMO), which weakens the immune response, improves viral replication, and may contribute to long COVID.

## Finding NEMO in the evolution of SARS CoV-2

The SARS-CoV-2 virus appears to have spilled over into humans relatively recently ^2^. Since the beginning of the SARS-CoV-2 pandemic, tremendous resources have gone into surveillance and sequencing worldwide, providing a detailed view of the evolution of this virus in its new human host, a process that continues to this day ^3^.

Two-thirds of the SARS-CoV-2 genome encodes the polyprotein pp1ab. This long precursor protein is processed by the papain-like protease (PL^pro^) and the

3-chymotrypsin-like protease, (3CL^pro^; also known as M^pro^ and NSP5) producing 16 nonstructural proteins (Nsp1-16) ^4–6^.

The conservation and function of these proteases make them excellent drug targets, ^7^ and Nirmatrelvir, an inhibitor of SARS-CoV-2 3CL^pr0^, was granted emergency use authorization in 2021 ^8^. 3CL^pro^ inhibition prevents the polyprotein processing essential for viral replication ^9^.

Surprisingly, in the emergency use authorization filed by Pfizer, it was noted that certain mutations in NSP5 arose in Nirmatrelvir-treated clinical trial patients ^8^. Some of the reported mutations map to the S1-S4’ subsites, surfaces adjacent to the catalytic site of the enzyme ^10^. Further, experiments in culture, where the virus is serially passaged in the presence of Nirmatrelvir, also produced similar mutations in the protease which rendered the enzyme insensitive to Nirmatrelvir inhibition ^8,11,12^. Surveys of the GSAID databases reveal that analogous mutations are in circulation ^13^. To date, the commonly accepted explanation for the appearance of these mutations in 3CL^pro^ has been that they arose due to the selection pressure of Nirmatrelvir, however, a different mechanism seems to be in play.

We have developed live cell assays for the main protease (3CL^pro^) activity. Cotransduction with two BacMam viruses delivers the protease and a fluorescent biosensor (3CLglow Up). In the assay, protease activity diminishes biosensor fluorescence, and the compounds that inhibit protease activity produce bright fluorescent cells. We developed the 3CLglowUp biosensor using the 8/9 cleavage site from the pp1ab SARS-CoV-2 protein. To develop assays for next-generation protease inhibitors, we created 20 mutant 3CL^pro^ proteases based on their reported Nirmatreliver insensitivity ^11–19^. Testing these mutant 3CL^pro^ proteases with our optimized 3CLglow Up sensor revealed that some mutants exhibited reduced activity compared to the parent WIV04 protease, while others showed improved performance.

Recent work has shown that the 3CL^pro^ also cleaves host immune signaling proteins, like nuclear factor-kB essential modulator (NEMO; also known as IKKy) ^20^. NEMO is necessary for activating the NF-κB pathway, a signaling pathway that rapidly responds to viral infection. The ablation of NEMO effectively dismantles the host immune response ^20,21^. Nemo cleavage by the main protease in infected brain endothelial cells leads to cell death and may contribute to the pathology of long COVID-19 ^22^. Additionally, NSP3, another non-structural protein in SARS-CoV-2, promotes the polyubiquination of NEMO/IKKy, disrupting NF-kB activation and modulating the host inflammatory response ^23^.

To investigate how the mutant proteases processed NEMO, we built an analogous protease sensor containing the NEMO cleavage site ERQ↓AREK. The results were surprising to us. All of the 3CL^pro^ mutants were as effective as, or more effective than, the 3CL^pro^ WIV04 strain at cleaving the NEMO-based protease sensor.

The sole driver of NSP5 evolution may be Nirmatrelvir. Our data, however, are consistent with a more nuanced model in which mutations are both conferring resistance to Nirmatrelvir while simultaneously increasing viral fitness to disarm the innate immune system.

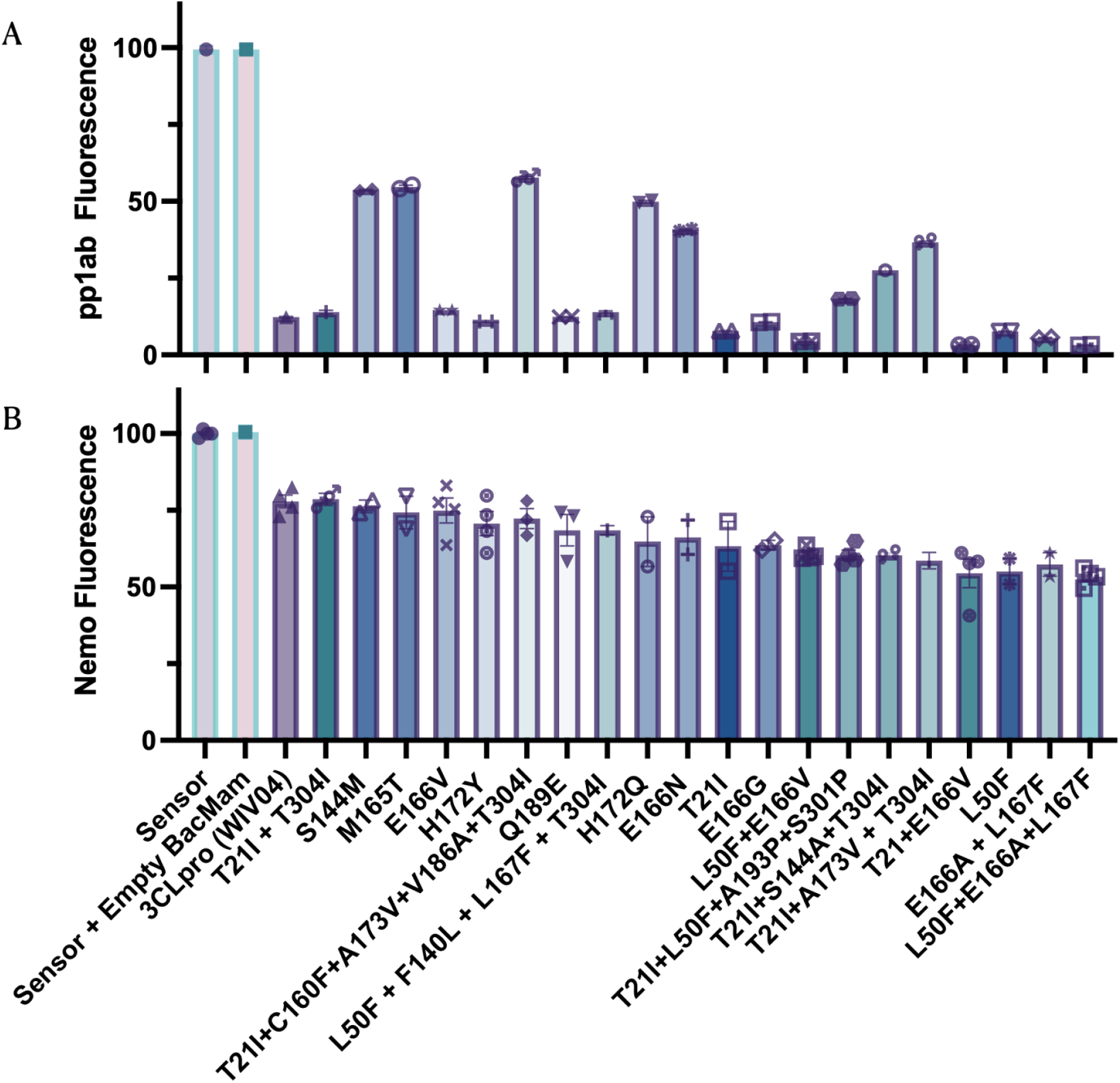
The protease assay involves co-transduction with two BacMam viruses that deliver the protease and a fluorescent biosensor. **(A)** HEK 293T cells were transduced with viruses expressing different mutant proteases, and the 3CLglow Up biosensor. The following day the cells were washed and fluorescence measured on a BioTek Neo plate reader. Some mutants cleaved the 3CLglow Up biosensor better than the parent enzyme, while others were significantly worse. **(B)** The same panel of proteases were tested against a biosensor carrying the NEMO cleavage site. The results were quite different, all of the mutants were as good as the parent enzyme or better at cleaving Nemo.

## Methods section

### Plasmids

The CMV expression plasmids were created using fragments synthesized by Integrated DNA Technologies (IDT, Coralville, IA, USA). Fragment assembly was performed using In-Fusion® HD Cloning Kit (Takara Bio USA) according to the manufacturer’s instructions. The expression plasmids of M^pro^ variants were made with PCR and primers designed to introduce the relevant mutations. We used the official reference sequence Wuhan WIV04 2019 from the Global Initiative on Sharing Influenza Data (GISAID) which is representative and identical to early outbreak sequences. The WIV04 strain was isolated by the Wuhan Institute of Virology from a clinical sample in 2019. The biosensor based on the 8/9 cleavage site used 213 amino acids surrounding the site, while the biosensor based on the NEMO cleavage site used 315 amino acids. All constructs were subjected to sequencing to verify coding regions using Eurofins Genomics EZ plasmid. The plasmids were then inserted into the BacMam genome through transposition ^1^, and the virus was produced in Sf9 cells and quantified by qPCR.

### Cell Line

HEK293T cells (human embryonic kidney; ATCC, VA, USA) were maintained in a growth medium composed of Dulbecco’s Modified Eagle Medium (DMEM, Life Technologies) supplemented with 100U/mL penicillin (Life Technologies), and 10% FBS (Signma-Aldrich, St. Louis, MI, USA) at 37°C in 5% CO_2_.

### Transduction

HEK293T cells were transduced with BacMam. All BacMam was standardized to 2x10^10^ viral genes per mL. The transduction mixture, sodium butyrate, and cells were plated at a cell density of 40,000 cells per well in 96-well or 384-well plates and left in the hood for 45 minutes. Following initial incubation in the hood, they are moved into the incubator. Twenty-four hours later the media was removed and replaced with DPBS. The plates were then read using the BioTek Synergy MX taking an endpoint fluorescence read with excitation/emission at 488/525 respectively.

